# Melanin concentrating hormone projections to the nucleus accumbens enhance the reward value of food consumption and do not induce feeding or REM sleep

**DOI:** 10.1101/2024.11.11.622987

**Authors:** Katherine L. Furman, Lorelei Baron, Hannah C. Lyons, Timothy Cha, Jack R. Evans, Jayeeta Manna, Limei Zhu, Joanna Mattis, Christian R. Burgess

**Affiliations:** Michigan Neuroscience Institute, University of Michigan, Ann Arbor, MI USA; Neuroscience Graduate Program, University of Michigan, Ann Arbor, MI USA; Department of Molecular and Integrative Physiology, University of Michigan, Ann Arbor, MI USA; Department of Neurology, University of Michigan, Ann Arbor, MI USA

**Keywords:** Melanin-concentrating hormone, Nucleus accumbens, Feeding, REM sleep, Reward, Lateral hypothalamus

## Abstract

Regulation of food intake and energy balance is critical to survival. Hunger develops as a response to energy deficit and drives food-seeking and consumption. However, motivations to eat are varied in nature, and promoted by factors other than energy deficit. When dysregulated, non-homeostatic drives to consume can contribute to disorders of food intake, adding to the increasing prevalence of restrictive eating disorders and obesity. Melanin-concentrating hormone (MCH) neurons have been implicated in the regulation of feeding behavior, in addition to a number of other fundamental behaviors including sleep, anxiety, and maternal behavior. Several studies suggest that MCH peptide increases food consumption, while studies of MCH neurons show effects only on cued feeding, and others show no effect of MCH neuron manipulation on feeding. MCH neurons have widespread projections to diverse downstream brain regions yet few studies have investigated the function of specific projections or differentiated the behaviors they regulate. Here we use optogenetics, in combination with different behavioral paradigms, to elucidate the role of MCH projections to the nucleus accumbens (NAc) in sleep and feeding behavior. We show that MCH neurons projecting to the NAc do not induce changes in baseline feeding or REM sleep, but do enhance the preference for a food paired with optogenetic stimulation. Furthermore, this effect is diminished in female mice relative to males, in line with previous results suggesting sex differences in the functional role of MCH neurons. These results suggest that MCH projections to the NAc can enhance the rewarding value of consumed food.

**Significance Statement:** While feeding is often driven by hunger, there are non-homeostatic reasons why animals consume food. Melanin-concentrating hormone (MCH) neurons have been implicated in the regulation of many fundamental behaviors, including feeding, sleep and reward. They project broadly throughout the brain, suggesting that they may mediate this diverse set of behaviors independently via specific projections to downstream regions. We used optogenetic activation of MCH neurons and their projections to the nucleus accumbens (NAc) in combination with complex behavioral paradigms to demonstrate that MCH projections to the NAc do not induce baseline feeding or increases in REM sleep but do enhance the value of a paired food. These results suggest that MCH neurons contribute to non-homeostatic consumption via projections to the NAc.

## Introduction

Animals must make informed decisions about which foods to eat, and how much, to maintain energy balance. Yet homeostatic need is not the sole factor in the decision to consume food. Humans commonly experience non-homeostatic motivators to eat, such as craving of sugary or fatty foods even when sated (i.e., “there is always room for dessert”). Over the last several decades there has been a steady increase in the prevalence of eating disorders and obesity in the United States, and these conditions are exacerbated by non-homeostatic feeding (Sturm and Hattori, 2013; Imes and Burke, 2014; Hoek, 2016; Ward et al., 2019; Cawley et al., 2021; Liu et al., 2021; Streatfeild et al., 2021). The specific mechanisms and circuitry through which the brain regulates non-homeostatic food intake are not fully understood.

The melanin-concentrating hormone (MCH) system has been implicated in homeostatic and non-homeostatic motivators to eat, in addition to a number of other fundamental behaviors. Knockout of Pmch from neurons in the lateral hypothalamus (LH) and zona incerta (ZI) results in mice that are hypophagic and lean, while overexpression or injection of MCH peptide into the brain induces feeding and can induce weight gain (Rossi et al., 1997; Shimada et al., 1998; Ludwig et al., 2001; Georgescu et al., 2005; Guesdon et al., 2009). MCH KO mice have shown normal overeating behavior in response to food restriction, a homeostatic motivator, but do not overeat in response to non-homeostatic conditioned food cues (Sherwood et al., 2015). More recent applications of optogenetics or pharmacogenetics have had mixed effects; stimulation or inhibition of MCH neurons shows no increase in feeding behavior (Dilsiz et al., 2019), or small effects under specific conditions (Terrill et al., 2020). Optogenetic activation of MCH neurons is reinforcing and, when explicitly paired with food availability, can increase consumption (Dilsiz et al., 2019). These data suggest a complex relationship between the MCH system and feeding, where specific projections and products of MCH neurons may drive homeostatic or non-homeostatic consumption.

MCH neurons have broad projections throughout the brain and are also implicated in the regulation of many other fundamental behaviors, including reward, reproduction, learning, anxiety, and sleep (Chaki et al., 2005; Potter and Burgess, 2022; Beekly et al., 2023; Concetti et al., 2024). As most investigations into the MCH system have focused only on one behavioral readout at a time and treated the population as homogeneous, it is unclear how the MCH system regulates non-homeostatic feeding and whether this is independent of effects on homeostatic feeding and other behaviors, like sleep. Here, we use optogenetic activation of both MCH neurons and their terminals in the nucleus accumbens (NAc) to demonstrate that MCH stimulation does not enhance food intake per se but enhances the rewarding value of consummatory behavior via projections to the NAc. Furthermore, stimulation of MCH terminals in the NAc does not elicit REM sleep, while stimulation of MCH neurons in the LH does. These data provide insights into the role of the MCH system in consummatory behavior and demonstrate the utility of investigating specific subpopulations and/or efferent projections of MCH neurons to more comprehensively elucidate their function across disparate behaviors.

## Methods

### Mice

All mice were housed in a University of Michigan vivarium in a temperature-controlled environment (12 h light and 12 h dark cycle; lights on at 2 AM) with *ad libitum* access to food and water. All animal protocols were approved by the University of Michigan’s Institutional Animal Care and Use Committee and are in accordance with NIH guidelines for the use and care of laboratory mice. For behavioral experiments, Pmch- ChR2-eYFP mice (n = 60) were made by crossing Ai32 mice (Strain #: 024109) to Pmch-Cre (RRID: IMSR_JAX:014099). Control mice were Cre-positive, ChR2-negative littermates. For rabies tracing experiments, Pmch-iCre positive mice (n = 5) were used (RRID: IMSR_JAX:014099, (Beekly et al. 2020)). Both male and female mice were used in all experiments. Overall health of experimental mice was monitored daily and any mice that displayed obvious signs of distress or weakness were removed from the study.

### Surgery

Mice were deeply anesthetized by inhalation of 2% isoflurane and placed on a Kopf stereotaxic apparatus (Tujunga, CA). Following standard disinfection procedure, the skin was retracted to expose the skull and a small hole was drilled into the skull unilaterally at defined positions to target NAc or LH. An optic fiber (400-µm diameter core; NA 0.5; RWD) was implanted unilaterally over NAc (A/P: 1.0 mm, M/L: −0.8 mm, D/V: - 4.2 mm) or LH (A/P: −1.4 mm, M/L: −1.0 mm, D/V: −4.7 mm). Fibers were fixed to the skull using dental acrylic. A subset of mice were also implanted with two stainless-steel screws (Frontal: A/P: 1.5 mm, M/L: −1.5 mm; Temporal: A/P: −3.5 mm, M/L: −2.8 mm) and a pair of multi strand stainless steel wires inserted into the neck extensor muscles. EEG and EMG leads were wired to a small electrical connector that is attached to the skull with dental cement.

Following surgery, mice were kept on a warming pad until fully ambulatory and fed a regular chow diet throughout the experimental period unless otherwise noted. All mice were given analgesics (5 mg/kg Meloxicam) prior to the end of surgery and 24 hours after surgery. Mice were given a minimum of 2 weeks for recovery before being used in any experiments. After the completion of the experiments, mice were euthanized and the locations of optic fiber tips were verified histologically.

For rabies tracing experiments, *Pmch-iCre* positive mice underwent an initial round of injections following anesthesia induction with 2-4% isoflurane. Mice were placed in a stereotaxic apparatus (KOPF Model 963). After exposing the skull via a small incision, a hole was drilled for injection. A pulled glass pipette (∼20 μm tip diameter, Friedrich & Dimmock, Inc.) was inserted into the brain, and AAV-hSyn-DIO-TVA(66T)- tdTomato was injected using a picospritzer. The pipette was kept in place for 5 minutes before withdrawal. Injections targeted the LH (A/P: −1.6 mm, M/L: −1.0 mm, D/V: −5.4 mm and −5.0 mm) and ZI (A/P: −1.6 mm, M/L: −1.0 mm, D/V: −4.6 mm and −4.3 mm). Postoperative care included intraperitoneal meloxicam (5 mg/kg) prior to the end of surgery and 24 h after surgery, as well as close monitoring for ten days to ensure recovery. Four to eight weeks later, 50-60 nL of rabies virus deltaG-N2c-Rabies-GFP (1x10^8 vg/mL) was injected into the ipsilateral nucleus accumbens shell (NAcSh; A/P: +1 mm, M/L: −0.52 mm, D/V: −4.6 mm). Mice were sacrificed five to eight days post-rabies injection.

## Experimental Design and Statistical Analysis

### Feeding and sleep experiments

Behavioral experiments began no less than 10 days following surgery. Mice were outfitted with an EEG/EMG tether in their home cage. To give mice time to habituate to the attached EEG/EMG tether, no optogenetic stimulation was delivered during a habituation phase which lasted at least 4 days. Once mice were adequately habituated to the tether (ambulatory, eating and drinking normally) optogenetic stimulation was delivered (20Hz, 10ms pulse width, 1s-ON 4s-OFF) for 3 hours during either the light cycle or the dark cycle, with alternating days of stimulation or baseline recording. Polysomnographic signals were digitized at 1000 Hz, with a 0.3-100 Hz bandpass filter applied to the EEG and a 30-100 Hz bandpass filter applied to the EMG (ProAmp-8, CWE Inc.), with a National Instruments data acquisition card and collected using a custom MATLAB script. EEG/EMG signals were notch filtered at 60 Hz to account for electrical interference from the recording tether. Polysomnographic signals were analyzed using AccuSleep, an open-source sleep scoring algorithm in MATLAB (Barger et al., 2019), and verified by an experienced sleep-scorer blinded to the group/condition. Behavioral states were scored in 5 s epochs as either Wake, non-rapid eye movement (NREM) sleep, or rapid eye movement (REM) sleep.

Animals’ home cages were modified to include an automatic feeder (FED3; (Matikainen-Ankney et al., 2021)). The FED3 was programmed to release a new food pellet (BioServ dustless precision pellets, 20mg) each time the previous pellet was taken by the mouse, such that food was always available. Feeding was quantified as the number of pellets consumed. Comparisons were made between days with and without optogenetic stimulation, as well as between control and ChR2 experimental groups. Sleep behavior was quantified by time spent in each arousal state, number of bouts of each arousal state, and lengths of bouts of each arousal state with a focus on REM sleep.

### Optogenetics-Reinforced Consumption Assay

Animals’ home cages were outfitted with two FED3 feeding devices (Matikainen-Ankney et al., 2021).

FED3s were programmed to release a food pellet (BioServ Dustless Precision Pellets, 20mg) following appropriate nose pokes. In order to pair optogenetic stimulation and pellet consumption, FED3s also acted as a closed-loop optogenetics trigger, activating optogenetic stimulation (20Hz, 10ms pulse width for 15s) upon appropriate pokes in the nose ports. While in the modified home cage, mice could move freely while attached to an optic fiber patch cable.

This paradigm has three phases: Habituation, StimON, and Reversal. The four-day Habituation phase allows mice to habituate to the optic fiber (day 1) and demonstrate any baseline preference among food ports (days 2-4). During Habituation, a poke in any of the four ports results in the same outcome: food pellet delivery. During the StimON phase, the FED3s were programmed to allow all-pairwise combinations of food and optogenetic stimulation at the four different ports. Ports were identified as (1) a Pellet + Stim port, which triggers both pellet delivery and optogenetic stimulation (20Hz for 15s), (2) a Stimulation Only port, which triggers only optogenetic stimulation (20Hz for 15s), (3) a Pellet Only port, which triggers only food pellet delivery, and (4) a No Outcome port, which does not result in either stimulation or pellet delivery. The StimON phase of the experiment lasted 5 days, a length we experimentally validated as it is long enough for the mouse to develop and demonstrate consistent preference. The last phase of the experiment is the Reversal phase, which is one day immediately following the end of the StimON phase. During Reversal, all four ports delivered a food pellet in response to a poke, and none triggered stimulation. In the results and figures, Sham stimulation refers to those opportunities where a mouse could nose poke to a port that either *will later* (Habituation phase) or *had previously* (Reversal phase) been paired with optogenetic stimulation, during these phases the LED was powered off.

The FED3 data collection includes data regarding both quantity and timing of pokes at each port. Using this data, the number of pokes in each of the four ports was counted on each day, and port preference was calculated as a percentage of total pokes that day. We also collected data of the cumulative pokes over time, to measure changes in port preference within-days by comparing the rate of increase across shorter durations.

### Preference Index

Mouse poking behavior at rewarded ports was used to calculate a preference index, indicating their preference for the Pellet+Stim port over other available ports. The calculation was as follows:

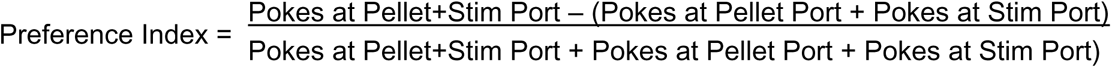

Preference index was calculated per-animal per-day, and mean preference index was compared across days and groups as a readout of their Pellet+Stim preference.

### Feeding Architecture

In addition to raw numbers of pokes and pellets, animals’ feeding architecture was quantified. This involved measuring inter-pellet intervals (IPIs) to understand the distribution of consumption over time. A histogram of IPI lengths was created to understand the distribution of IPI lengths for a given animal. This histogram was used to determine the appropriate interbout interval, in order to denote separate feeding “bouts.” A majority of IPIs were less than 2 minutes, thus 2 minutes was determined as the interbout interval. Using these criteria, the number of feeding bouts and the number of food pellets per bout were calculated. The skewness of the IPI distribution was also calculated.

### Acute slice preparation

Mice were deeply anesthetized using inhaled isoflurane. Brains were removed and then immediately placed in an ice-cold sucrose solution containing (in mM): 87 NaCl, 75 sucrose, 2.5 KCl, 1.0 CaCl2, 2.0 MgSO4, 26 NaHCO3, 1.25 NaH2PO4, and 10 glucose, maintained with 95% O2 and 5% CO2. Using a Leica VT-1200S vibratome (Leica Microsystems Inc., Buffalo Grove, IL, USA), coronal slices of 300 μm thickness were prepared. These slices were then transferred to a holding chamber filled with oxygenated artificial cerebrospinal fluid (ACSF) comprised of (in mM): 125 NaCl, 2.5 KCl, 2.0 CaCl2, 1.0 MgSO4, 26 NaHCO3, 1.25 NaH2PO4, and 10 glucose. The slices were allowed to recover at 32°C for 30 minutes, followed by an additional 30 minutes at room temperature before proceeding to recordings.

### Slice electrophysiology

Whole-cell patch-clamp recordings were conducted using a SliceScope Pro 6000 electrophysiology system (Scientifica). Slices were transferred to a recording chamber and continuously perfused with oxygenated ACSF at a flow rate of roughly 3 mL/min. All slice physiology experiments were carried out at 31°C. MCH neurons were visualized using a SliceScope microscope (Scientifica) and identified by their YFP expression. Borosilicate glass electrodes, created using a P-97 puller (Sutter Instruments) to achieve a tip resistance of 3-4 MΩ, were used for whole-cell voltage- and current-clamp recordings. Electrodes were filled with a K-Gluconate internal solution, composed of (in mM): 130 K-gluconate, 6.3 KCl, 0.5 EGTA, 1.0 MgCl2, 10 HEPES, 4.0 Mg-ATP, and 0.3 Na-GTP. The pH was adjusted to 7.30 with KOH and the osmolarity was calibrated to 285 mOsm with 30% sucrose. Electrodes were maneuvered using a PatchStar manipulator (Scientifica). Signals were sampled at 20 kHz, amplified using a MultiClamp 700B amplifier (Molecular Devices), filtered at 10 kHz, digitized with a DigiData 1550, and captured using pClamp10 software. Data were included in the final analysis only if the cell had a resting potential below −50 mV. Series resistance compensation (bridge balance) was applied throughout the current clamp experiments, with readjustments made as necessary. No correction for liquid junction potential was performed. Activation of ChR2 was achieved through widefield blue (470 nm) LED illumination (CoolLED pE-800; Scientifica).

### Immunohistochemistry

Following the conclusion of all experiments, mice were anesthetized with pentobarbital and transcardially perfused with phosphate-buffered saline (PBS) followed by formalin. Once removed, brains were post-fixed in 10% formalin overnight then transferred to 20% PBS-sucrose for ∼48 hours. Brains were then frozen and 30μm coronal sections were cut using a freezing microtome. Location of optic fibers was confirmed by histology, and mice in which the fiber did not hit the intended target area were excluded.

For MCH immunohistochemistry, coronal sections were washed with 1x PBS and then incubated in 10% NDS (Normal Donkey Serum) blocking solution for 1 hour at room temperature. Once blocked, sections were incubated in MCH primary antibody solution (Rabbit anti-MCH Polyclonal Antibody (Catalog #H-070-47, Phoenix Pharmaceuticals) (1:8000)) overnight at RT. Following primary antibody staining, sections were washed with PBS-Tween then incubated in secondary antibody solution (Donkey anti-Rabbit Alexa Fluor 647 Conjugated Antibody (Catalog #A-31573, Invitrogen) (1:1250)) for 1 hour at RT. Slices went through a final PBS wash and then mounted onto glass slides using Vectashield Antifade Mounting Medium with DAPI (Vector Laboratories, Burlingame, CA). Images were then captured by a fluorescence microscope (Olympus IX83/Keyence BZ-X800) at 10x magnification.

To increase the brightness of endogenous eYFP fluorescence, we used GFP amplification. Sections were washed with PBS then incubated in 5% NDS blocking solution for 2 hours at RT. After blocking, sections were incubated in primary antibody solution (GFP Polyclonal Antibody, Alexa Fluor™ 488 (Catalog #A-21311, ThermoFisher) (1:1000)) for 48 hours at RT. After primary antibody incubation, sections were washed with PBS then mounted onto slides.

For rabies tracing experiments, brain slices in a 1-in-3 series were mounted on slides and cover- slipped. Images were taken using a BZ-X800 fluorescence microscope. Coronal fluorescent images were registered to the Allen Brain Atlas V3p1 using the Fiji plugin ABBA for accurate mapping. Once registered, images were transferred to Adobe Illustrator to align with the appropriate Allen Brain Atlas template, allowing simultaneous display of the entire brain slice and cell distribution. The fluorescent ABBA brain sections were used to mark all MCH neurons containing mCherry, GFP, or mCherry and GFP with a filled circle using the ellipse tool. The relative positions of these circles were mapped onto the Allen Brain Atlas template. Anything above the NS tract was considered ZI, anything at or below was considered LH (Watson et al., 2014).

Neurons were marked only if the soma was clearly visible.

## Statistical analysis

Statistical analyses were performed using Prism 9.0 (GraphPad), IBM SPSS Statistics, or Matlab software. Details of statistical tests employed can be found in the relevant figure legends. In all statistical tests, normal distribution and equal variance was established. The data presented met the assumptions of the statistical test employed. Exclusion criteria for experimental mice were (i) sickness or death during the testing period (ii) if histological validation of the injection site demonstrated an absence of reporter gene expression (iii) or if histological validation demonstrated a mistargeted fiber placement. These criteria were established before data collection. N numbers represent final numbers of healthy/validated mice.

## Results

We sought to elucidate the function of MCH neurons and their projections to the NAc by generating an MCH-ChR2 transgenic mouse (Figure 1A). To validate that MCH neurons were sensitive to our optogenetic stimulation parameters, ChR2-expressing MCH neurons were held in voltage clamp (−70 mV) and demonstrated photocurrent when exposed to 1 s pulses of blue light (470 nm, 14.5 mW/mm2; Figure 1B).

**Figure 1.**
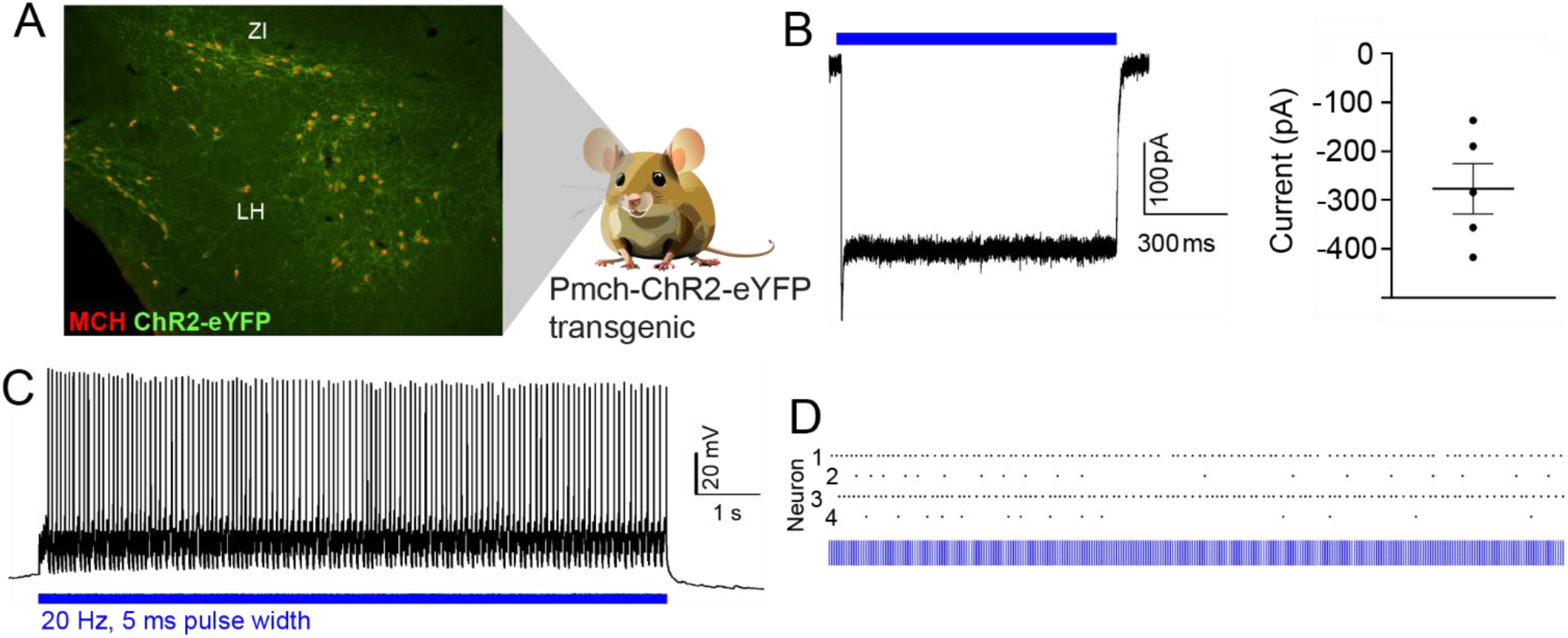
Optogenetic activation of MCH neurons in MCH-ChR2 mice. A. MCH-ChR2 mice express eYFP-labeled ChR2 in MCH neurons. B. ChR2-expressing MCH neurons were held in voltage clamp (−70 mV) and exposed to 1 s pulses of blue light (470 nm, 14.5 mW/mm2). Left: A representative photocurrent in an MCH neuron in response to light (blue horizontal bar). Right: Steady state photocurrents (−227 ± 52 pA; n = 5 cells). C. Representative current clamp trace from a ChR2-expressing MCH neuron in which a sustained train of action potentials was evoked via photostimulation (470 nm, 14.5 mW/mm2, 20 Hz, 5 ms pulse width). D. Action potentials (black dots) elicited by photostimulation in four MCH neurons.

Current clamp experiments demonstrated sustained trains of action potentials evoked via blue light photostimulation (470 nm, 14.5 mW/mm2, 20 Hz, 5 ms pulse width; Figure 1C,D). Not all recorded neurons were able to follow continuous 20 Hz stimulation (Figure 1D). These *ex vivo* slice electrophysiology recordings from MCH-ChR2 mice confirmed that MCH neurons were sensitive to light stimulation.

MCH neurons send dense projections to the NAc, in particular the medial NAc shell (Georgescu et al. 2005). We used rabies-based retrograde tracing to demonstrate the proportion and location of MCH neurons that send projections to the NAc (Figure 2A). Retrogradely labeled NAc-projecting neurons were prevalent in the MCH field (Figure 2B; n = 5 mice). Cell counts demonstrated that 21.4 +/- 2.2% of all MCH neurons send projections to the NAc (Figure 2B, inset). The majority of MCH neurons projecting to the NAc were located in the LH rather than the ZI (LH: 95.4 +/- 1.1% of NAc-projecting neurons originated in the LH; Figure 2C), with a higher proportion of LH neurons also projecting to NAc (LH: 24.2 +/- 2.9% of LH neurons project to the NAc; ZI: 10.2 +/- 2.5% of ZI neurons project to the NAc).

**Figure 2.**
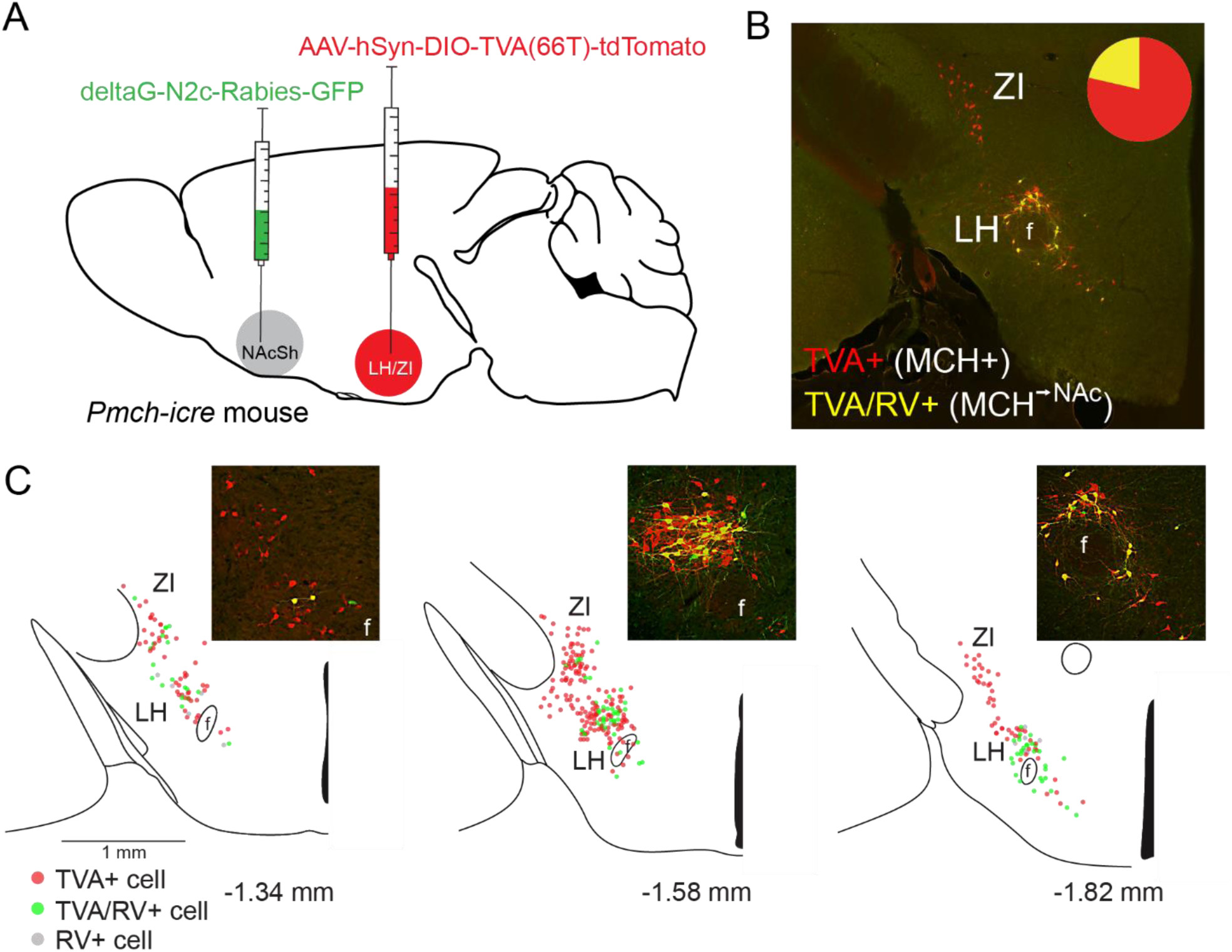
MCH neurons send dense projections to the NAc. A. Schematic of rabies-based retrograde tracing experiments. B. Example image demonstrating TVA-expressing MCH neurons (red) and retrogradely labeled rabies- expressing MCH neurons (yellow) that project to the NAc. 21.4 +/- 2.2% of MCH neurons project to the NAc (n = 5 mice; inset). C. Composite images showing the distribution of TVA-labeled neurons and rabies virus-labeled neurons in the lateral hypothalamus (LH) and zona incerta (ZI). 24.2 +/- 2.9% of LH MCH neurons project to the NAc while 10.2 +/- 2.5% of ZI MCH neurons project to the NAc. Insets show individual example images from the LH. f: fornix.

### MCH projections to the NAc do not modulate REM sleep or baseline feeding

We next aimed to understand the role of MCH neurons and their projections to the NAc on homeostatic behaviors; feeding and sleeping. We recorded polysomnographic signals with EEG/EMG tethers and feeding behavior using a FED3 food hopper (Matikainen-Ankney et al., 2021) as mice freely behaved in their homecage (Figure 3A). Mice had an optic fiber implanted in either the LH or the NAc, and received three hours of optogenetic stimulation (20Hz, 1s on, 4s off) during the light and dark cycle on alternating days. While mice expectedly consumed more food pellets during the dark cycle than the light cycle, the amount of food pellets consumed in each phase was similar regardless of whether they received optogenetic stimulation (Figure 3B,C). We did observe an effect of MCH optogenetic stimulation on REM sleep in LH, but not in NAc (Figure 3E,F). Representative hypnograms of an animal with the optic fiber in the LH are shown in Figure 3D. Animals with optic fibers placed in the LH exhibited a significant increase in REM sleep bout length during the light cycle on days with optogenetic stimulation compared to days without (paired t-test, t(16) = 2.589; p= 0.0198; Figure 3E, top). These mice also showed a trend toward an increase in the number of REM bouts during the dark cycle (paired t-test, t(16) =1.567; p= 0.1366; Figure 3E, bottom). No effect of MCH stimulation on REM sleep was observed for mice with optic fibers placed in the NAc (Figure 3F). There was also no effect of optogenetic stimulation on other arousal states (NREM or Wake; Data not shown).

**Figure 3.**
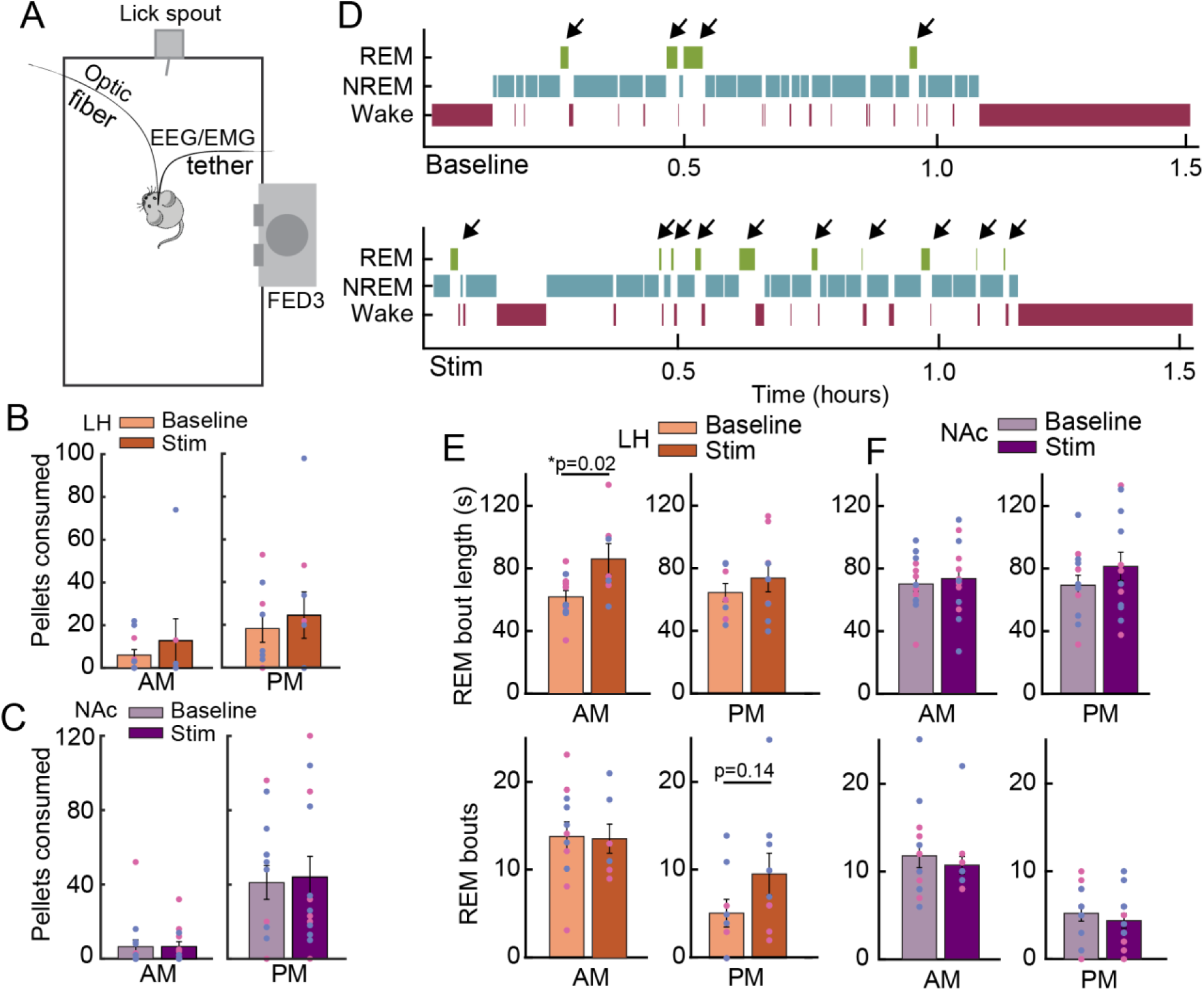
Continuous optogenetic stimulation of the MCH system does not increase food intake but can modulate REM sleep. A. Homecages were modified to include a Feeding Experimentation Device 3 (FED3) from which mice could retrieve food pellets ad libitum. Mice were outfitted with both an optic fiber to deliver optogenetic stimulation and an EEG/EMG recording tether to monitor sleep-wake architecture. Fiber implant locations are shown in Extended Data Figure 3-1. B. Optogenetic stimulation of MCH neurons in the LH did not increase food pellet intake during either the light or dark phase (paired t-test, p>0.05). Control mice with fiber in LH also show no change in feeding behavior (Extended data figure 3-1B). There was no significant change in meal architecture as a result of LH MCH optogenetic stimulation (Extended Data Figure 3-2A,B), and feeding behavior was similar for both males and females (Extended Data Figure 3-3A, right). Pink and blue dots denote female and male mice, respectively. C. Optogenetic stimulation of MCH terminals in the NAc did not increase food pellet intake during either the light or dark phase (paired t-test, p>0.05). Control mice with fiber in NAc also show no change in feeding behavior (Extended data figure 3-1C). There was no significant change in meal architecture as a result of MCH optogenetic stimulation (Extended Data Figure 3-2C,D), and feeding behavior was similar for both males and females (Extended Data Figure 3-3B, right). Pink and blue dots denote female and male mice, respectively. D. Example hypnograms taken during the dark cycle from a MCH-ChR2 mouse with optic fiber implanted into the LH with optogenetic stimulation off (top) or on (bottom). Arrows denote REM sleep bouts. E. Optogenetic stimulation of MCH neurons in the LH led to a significant increase in REM bout length during the light cycle (top, paired t-test, t(16) = 2.589; p= 0.0198), and a non-significant change in the number of REM sleep bouts during the dark cycle (bottom, paired t-test, t(16) =1.567; p= 0.1366). Both male and female mice had similar changes in REM sleep architecture as a result of MCH optogenetic stimulation (Extended Data Figure 3-3A). Pink and blue dots denote female and male mice, respectively. F. There was no significant difference in REM sleep architecture with optogenetic stimulation of MCH terminals in NAc (paired t-test, p>0.05). Both male and female mice had similar changes in REM sleep architecture as a result of MCH optogenetic stimulation (Extended Data Figure 3-3B). Pink and blue dots denote female and male mice, respectively.

### MCH projections to the NAc enhance the value of consummatory behavior

As there was no effect of MCH → NAc stimulation on either acute feeding or arousal, we developed an optogenetics-reinforced consumption assay (ORCA) to assess how consumption behavior changes when mice have voluntary access to self-stimulation of MCH in LH, or MCH terminals in NAc. Following surgical implantation of an optic fiber in either LH or NAc, MCH-ChR2 mice were given the choice between all-pairwise combinations of food and optogenetic stimulation between two automated food hoppers (FED3) and four total ports. During the Habituation phase of the ORCA paradigm, when all ports resulted in delivery of a pellet and none resulted in optogenetic stimulation (Sham stimulation refers to a port that *will later* be paired with optogenetic stimulation), mice from all groups showed no significant innate preference for any individual port (two-way ANOVA/Bonferroni multiple-comparison test; LH group: stim, p = 0.165; F(1,16) = 2.144; pellet, p = 0.4715; F(1,16) = 0.5438; interaction, p = 0.7212; F(1,16) = 0.1320; NAc group: stim, p = 0.9133; F(1,16) = 0.01224; pellet, p = 0.8320; F(1,16) = 0.04548; interaction, p = 0.7447; F(1,16) = 0.1097) (Figure 4B,C, left).

**Figure 4.**
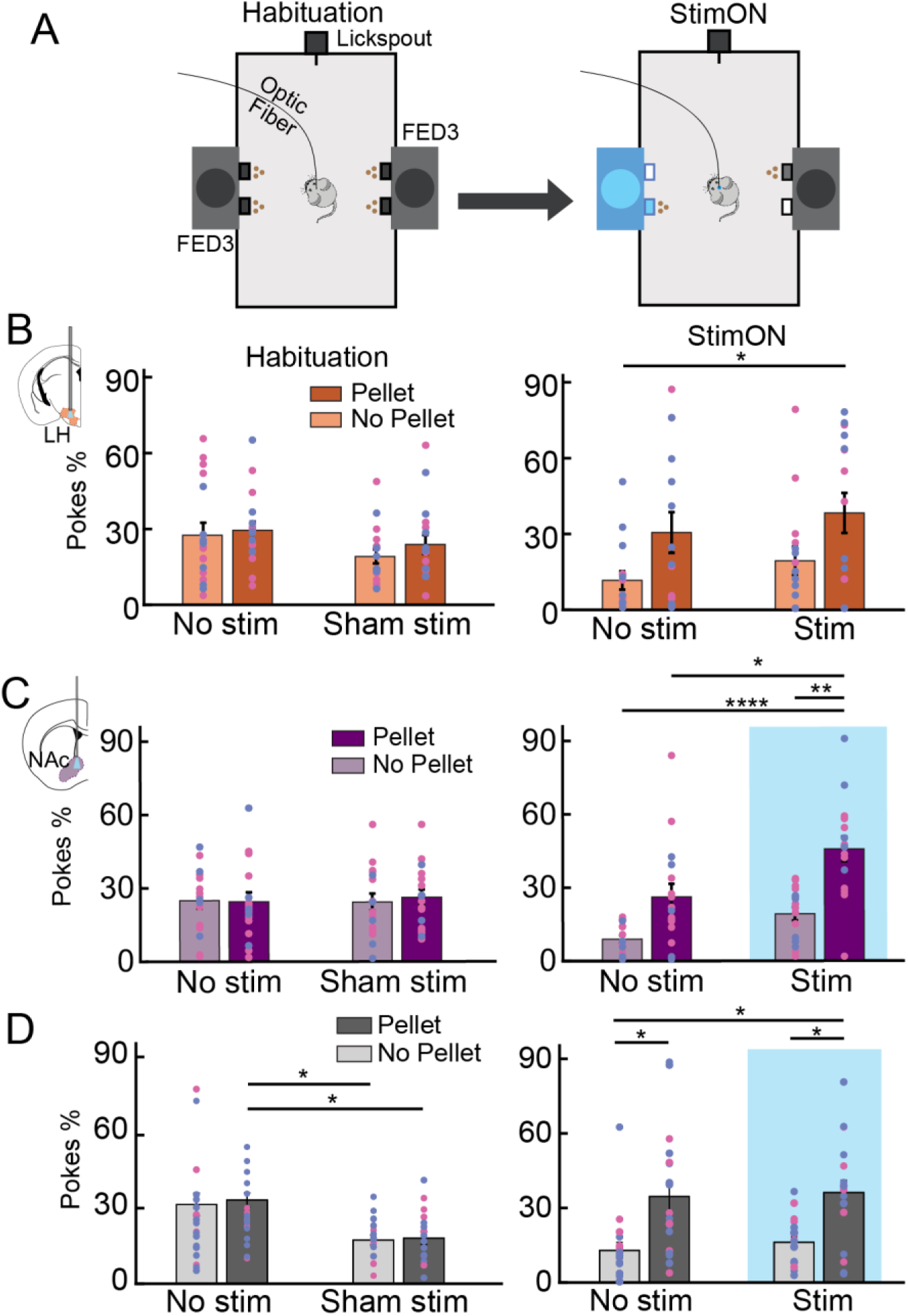
Optogenetic stimulation of MCH terminals in the NAc increases preference for a paired food. A. Optogenetics Reinforced Consumption Assay (ORCA) behavioral schematic. Mouse homecages were modified to include two FED3 food hoppers. During habituation, a poke at either nose port on each FED3 triggers pellet delivery from the pellet well. During StimON, pokes at one nose port on each FED3 trigger food pellet delivery, while pokes at both nose ports on one FED3 trigger optogenetic stimulation. B. Poke preference data from mice with a fiber implanted into the LH on the final Habituation day (left) and the final StimON day (right). There was no significant difference observed in port preference on the Habituation day (two-way ANOVA/Bonferroni multiple-comparison test; stim, p = 0.165; F(1,16) = 2.144; pellet, p = 0.4715; F(1,16) = 0.5438; interaction, p = 0.7212; F(1,16) = 0.1320). On the final StimON day, there was a significant effect of Pellet availability on port preference (two-way ANOVA/Bonferroni multiple-comparison test; stim, p = 0.4742; F(1,14) = 0.5409; pellet, **p = 0.0023; F(1,14) = 16.85; interaction, p = 0.9978; F(1,14) = 8.226e-006) Specifically, animals’ preference for the Pellet+Stim port was significantly higher than for the No Pellet+No Stim port (Bonferroni multiple-comparison test, p=0.0390). There were no significant changes to feeding architecture (Extended data figure 4-1, orange bars). Pink and blue dots denote female and male mice, respectively. C. Poke preference data from mice with a fiber implanted into the NAc on the final Habituation day (left) and the final StimON day (right). There were no significant differences observed on the Habituation day (two-way ANOVA/Bonferroni multiple-comparison test; stim, p = 0.9133; F(1,16) = 0.01224; pellet, p = 0.8320; F(1,16) = 0.04548; interaction, p = 0.7447; F(1,16) = 0.1097). On the StimON day (left), there was a significant effect of both stimulation and pellet availability on poke preference (two-way ANOVA/Bonferroni multiple-comparison test; stim, *p = 0.0332; F(1,15) = 5.498; pellet, ****p <0.0001; F(1,15) = 64.17; interaction, p = 0.2656; F(1,15) = 1.338). Mice receiving MCH optogenetic stimulation in the NAc significantly preferred the Pellet+Stim port over the Stim Only port (Bonferroni multiple-comparisons test, p= 0.0018), the Pellet Only port (Bonferroni multiple-comparisons test, p= 0.0217), and the No Pellet+No Stim port (Bonferroni multiple-comparisons test, p<0.001).There were no significant changes to feeding architecture (Extended data figure 4-1, purple bars). Pink and blue dots denote female and male mice, respectively. D. Poke preference data from control mice on the final Habituation day (left) and the final StimON day (right). Control mice showed a significant preference on the Habituation day (two-way ANOVA/Bonferroni multiple- comparison test; stim, ***p = 0.0003; F(1,17) = 20.51; pellet, p = 0.7563; F(1,17) = 0.09945; interaction, p = 0.8939; F(1,17) = 0.018939), and no significant preference on the final StimON day (two-way ANOVA/Bonferroni multiple-comparison test; stim, p = 0.6869; F(1,17) = 0.1681; pellet, ****p < 0.0001; F(1,17) = 27.35; interaction, p = 0.8543; F(1,17) = 0.03477). There were no significant differences based on fiber location in Control animals (Extended data figure 4-2). Pink and blue dots denote female and male mice, respectively.

Following Habituation, the ports delivering stimulation were assigned to the food hopper which the mouse had least preferred during habituation. This accounts for the slight preference Control mice (the MCH-Cre littermates of the MCH-ChR2 experimental mice) had for the two “No Stimulation” ports during their Habituation phase (two-way ANOVA/Bonferroni multiple-comparison test; stim, ***p = 0.0003; F(1,17) = 20.51; pellet, p = 0.7563; F(1,17) = 0.09945; interaction, p = 0.8939; F(1,17) = 0.018939) (Figure 4D, left). After observing no innate preference for any of the four ports, the paradigm was changed to allow mice a choice between food, stimulation, or both.

During the StimON phase of the ORCA paradigm, mice had access to all-pairwise combinations of food pellet delivery and optogenetic stimulation. Both MCH-ChR2 mice with the fiber in the LH and controls showed a significant and expected shift towards a preference for the two ports delivering a food pellet, and away from the ports that no longer deliver food pellets (two-way ANOVA/Bonferroni multiple-comparison test; stim, p = 0.4742; F(1,14) = 0.5409; pellet, **p = 0.0023; F(1,14) = 16.85; interaction, p = 0.9978; F(1,14) = 8.226e-006) (Figure 4B,D right; Extended data Figure 4-2). Specifically, animals’ preference for the Pellet+Stim port was significantly higher than for the No Pellet+No Stim port (Bonferroni multiple-comparison test, p=0.0390).

However, MCH-ChR2 mice with the fiber placed in the NAc showed a significant main effect of both food and stimulation availability on port preference (two-way ANOVA/Bonferroni multiple-comparison test; stim, *p = 0.0332; F(1,15) = 5.498; pellet, ****p <0.0001; F(1,15) = 64.17; interaction, p = 0.2656; F(1,15) = 1.338) (Figure 4C, right). Mice receiving MCH optogenetic stimulation in the NAc significantly preferred the Pellet+Stim port over the Stim Only port (Bonferroni multiple-comparisons test, p= 0.0018), the Pellet Only port (Bonferroni multiple-comparisons test, p= 0.0217), and the No Pellet+No Stim port (Bonferroni multiple-comparisons test, p<0.001). The preference for the port delivering both a food pellet and stimulation in the MCH-ChR2 group with MCH → NAc stimulation was not due to a change in the total number of pellets consumed, or a change in overall meal architecture (number of feeding bouts, or number of pellets per feeding bout; Extended data Figure 4-1) as, similar to homecage optogenetic stimulation, there was no increase in feeding or change in meal structure in the ORCA paradigm.

In addition to our investigation of port preference after each session, we also investigated the time course of nose poking and port preference emergence within ORCA sessions. A representative raster plot from a single mouse shows differences in poking patterns at each port (Figure 5A). For mice receiving MCH stimulation in the LH, both MCH-ChR2 and Control mice develop a persistent preference for ports with a food pellet, regardless of whether the food is paired with stimulation (Figure 5B, left) or not (Figure 5B, right).

**Figure 5.**
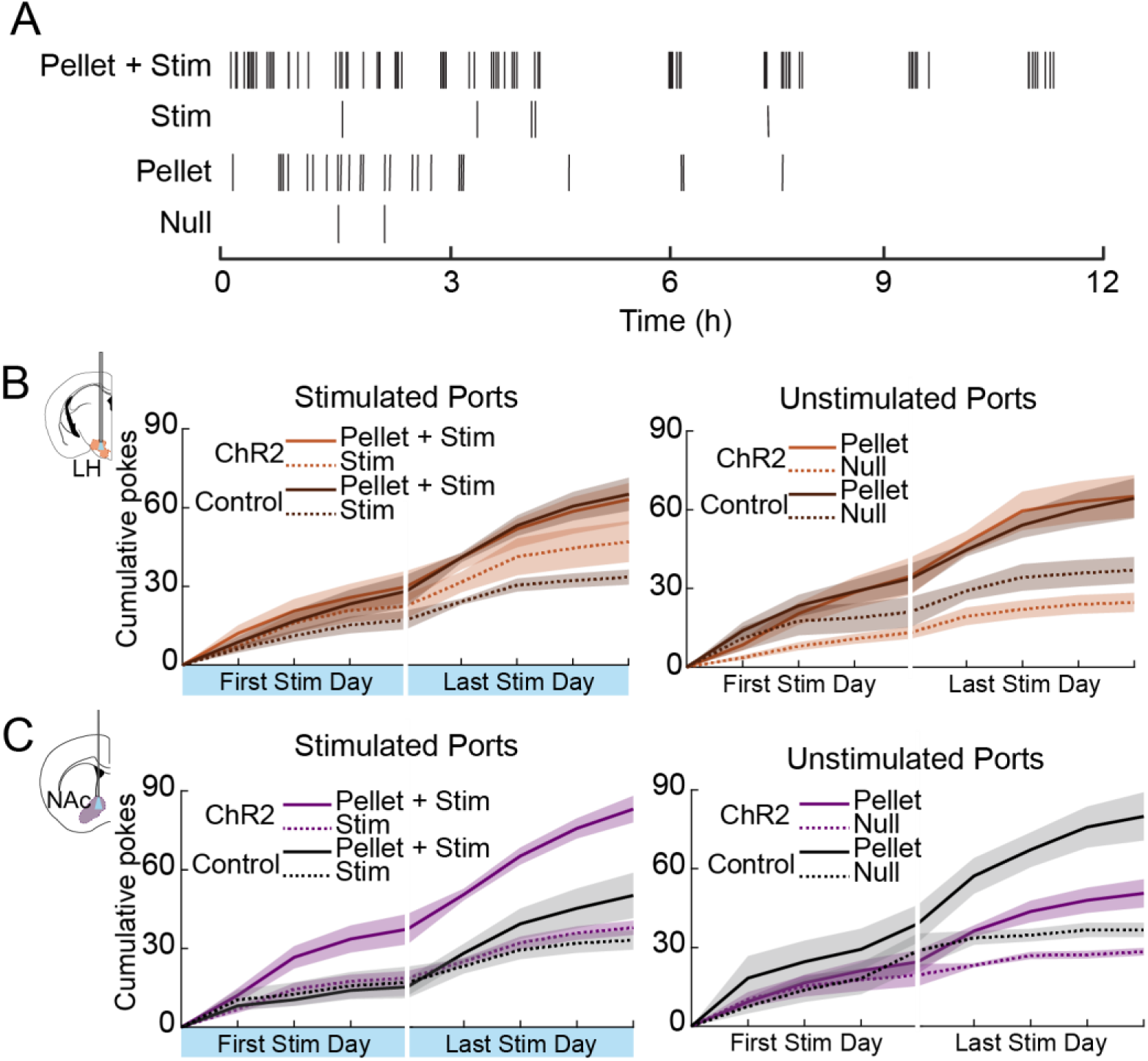
Preference for stimulation-paired food emerges more rapidly with MCH→ NAc optogenetic stimulation. A. Raster plot of a representative mouse’s poking behavior over the 12 hour dark cycle. Vertical lines represent pokes at the specified port. B. MCH-ChR2 mice optogenetic stimulation in LH show similar behavior to Control mice. The preference develops by the end of the final day of stimulation. C. MCH-ChR2 mice receiving MCH optogenetic stimulation in the NAc quickly develop a preference for the Pellet+Stim port, by the middle of the first day of stimulation, which lasts throughout the entire StimON period. Control mice receiving sham stimulation do not develop this preference, preferring the port(s) delivering a food pellet regardless of stimulation availability.

However, mice receiving MCH → NAc stimulation quickly develop a preference for the Pellet + Stim port, emerging during the first StimON dark period and lasting through the end of the StimON phase (Figure 5C, left). Control mice receiving sham stimulation in the NAc take longer to develop a preference, and only prefer pellet ports, regardless of whether the food pellet is paired with stimulation (Figure 5C).

The StimON phase of the ORCA experiment is followed by a day-long Reversal phase, during which the four ports all release a food pellet, and stimulation can no longer be triggered by poking (Sham stimulation refers to a port that *had previously* been paired with optogenetic stimulation). Following reversal of the paradigm, there remained a significant effect of prior food pellet availability in MCH-ChR2 mice receiving MCH optogenetic stimulation in LH (two-way ANOVA/Bonferroni multiple-comparison test; stim, p = 0.1933; F(1,16) = 1.844; pellet, *p = 0.0148; F(1,16) = 7.457; interaction, p = 0.7970; F(1,16) = 0.06838), and a significant effect of both prior stimulation and prior food pellet availability in MCH-ChR2 mice receiving MCH optogenetic stimulation in NAc. (two-way ANOVA/Bonferroni multiple-comparison test; stim, *p = 0.0189; F(1,13) = 7.178; pellet, *p = 0.0172; F(1,13) =7.458; interaction, p = 0.9396; F(1,13) = 0.005975; Figure 6 A,B). When the poking data of the Reversal phase are viewed cumulatively, it is clear that, regardless of fiber location, mice retain a preference for their previously-preferred port(s) during the first three hours of the Reversal phase (Figure C,D, left). However, this preference is entirely abolished by the end of the Reversal phase, as the poking behavior at each port diminishes, with mice failing to prefer any specific port during this time (Figure C,D, right).

**Figure 6.**
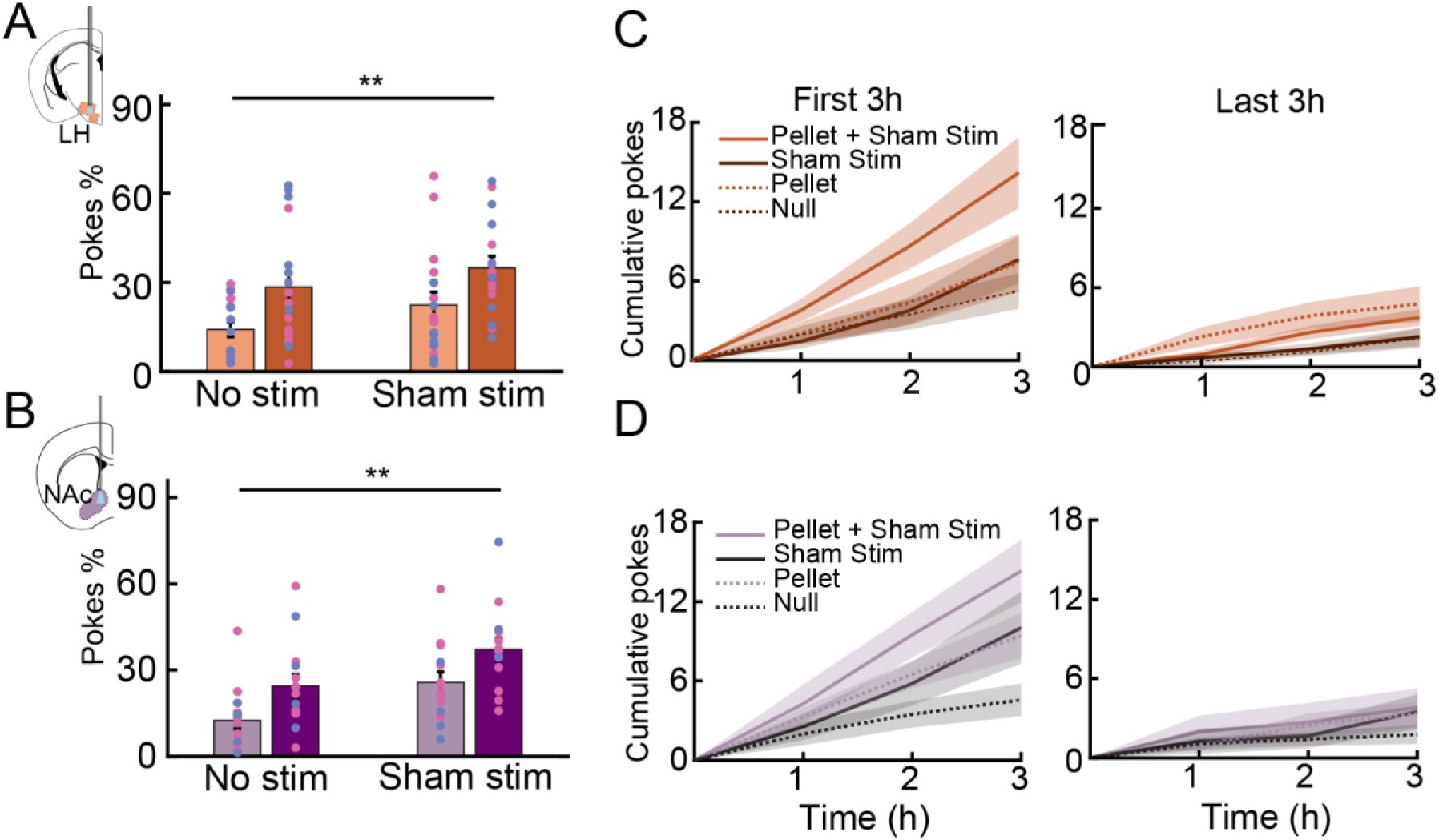
Preference persists for a short period after removal of stimulation. A. Percentage of pokes at each port during the Reversal day of the ORCA paradigm for mice receiving MCH optogenetic stimulation in LH. Following reversal of the paradigm, there remained a significant effect of food pellet availability (two-way ANOVA/Bonferroni multiple-comparison test; stim, p = 0.1933; F(1,16) = 1.844; pellet, *p = 0.0148; F(1,16) = 7.457; interaction, p = 0.7970; F(1,16) = 0.06838). Pink and blue dots denote female and male mice, respectively. B. Percentage of pokes at each port during the Reversal day of the ORCA paradigm for mice receiving MCH optogenetic stimulation in NAc. Following reversal of the paradigm, there remained a significant effect of both stimulation and food pellet availability (two-way ANOVA/Bonferroni multiple-comparison test; stim, *p = 0.0189; F(1,13) = 7.178; pellet, *p = 0.0172; F(1,13) =7.458; interaction, p = 0.9396; F(1,13) = 0.005975). Pink and blue dots denote female and male mice, respectively. C. Cumulative poking behavior over the first three hours (left) and last three hours (right) of the Reversal day of the ORCA paradigm for mice receiving MCH optogenetic stimulation in LH. A preference for pellet ports persisted during the first three hours of the Reversal day but was eliminated by the last three hours. D. Cumulative poking behavior over the first three hours (left) and last three hours (right) of the Reversal day of the ORCA paradigm for mice receiving MCH optogenetic stimulation in NAc. A preference for the Pellet+Stim port persisted during the first three hours of the Reversal day but was eliminated by the last three hours.

### Sex-specific effects of MCH stimulation

Some previous research has demonstrated sex-specific roles for the MCH system in feeding (Terrill et al. 2020; Ferreira et al. 2017). We sought to investigate whether there was a sex-specific effect of MCH optogenetic stimulation in our ORCA paradigm. During the StimON phase, effects of MCH optogenetic stimulation were more pronounced in males than females, (Figure 7), and this effect was stronger in the NAc. Females receiving optogenetic stimulation in LH (Figure 7A, top) showed no significant effect of either stimulation or food pellet availability (two-way ANOVA/Bonferroni multiple-comparison test; stim, p = 0.5837; F(1,5) = 0.3427; pellet, p = 0.3185; F(1,5) = 1.226; interaction, p = 0.2930; F(1,5) = 1.380). Males receiving optogenetic stimulation in LH (Figure 7A, bottom) showed a significant effect of food pellet availability (two-way ANOVA/Bonferroni multiple-comparison test; stim, p = 0.6750; F(1,8) = 0.1892; pellet, ***p = 0.0006; F(1,8) = 29.87; interaction, p = 0.2779; F(1,8) = 1.355). The pellet preference emerged over the course of the 12 h of stimulation (Figure 7B). We calculated a preference index to quantify the degree to which mice preferred the Pellet+Stim port relative to the other available ports. In this measure, male and female mice receiving LH MCH optogenetic stimulation did not have a significantly different preference for the Pellet+stim port (Figure 7C).

**Figure 7.**
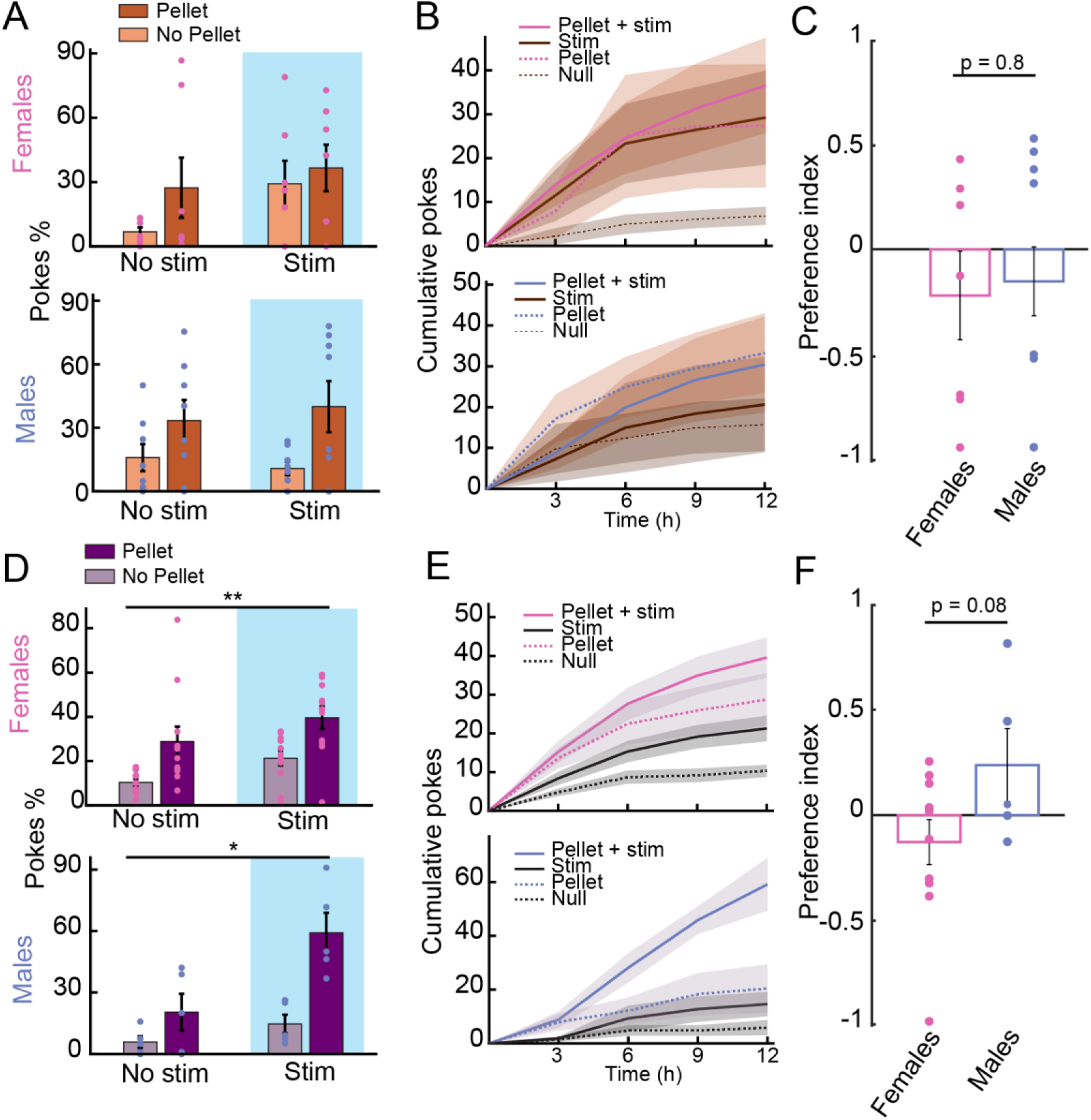
Sex-specific effects of MCH stimulation and associated preference. A. Poke percentages on the final day of the StimON phase of the ORCA paradigm for females (top) and males (bottom) receiving MCH optogenetic stimulation in LH. Females receiving optogenetic stimulation in LH (top) showed no significant effect of either stimulation or food pellet availability (two-way ANOVA/Bonferroni multiple-comparison test; stim, p = 0.5837; F(1,5) = 0.3427; pellet, p = 0.3185; F(1,5) = 1.226; interaction, p = 0.2930; F(1,5) = 1.380). Males receiving optogenetic stimulation in LH (bottom) showed a significant effect of food pellet availability (two-way ANOVA/Bonferroni multiple-comparison test; stim, p = 0.6750; F(1,8) = 0.1892; pellet, ***p = 0.0006; F(1,8) = 29.87; interaction, p = 0.2779; F(1,8) = 1.355). B. Cumulative poking behavior over the entire 12hr dark cycle on the final day of the StimON phase of the ORCA paradigm for females (top) and males (bottom) receiving MCH optogenetic stimulation in LH. No significant differences between ports were observed. Females receiving optogenetic stimulation in NAc (top) showed a significant effect of pellet availability (two-way ANOVA/Bonferroni multiple-comparison test; stim, p = 0.1856; F(1,10) = 2.021; pellet, ***p = 0.0001; F(1,10) = 35.00; interaction, p = 0.9888; F(1,10) = 0.0002062). Males receiving optogenetic stimulation in NAc(top) showed a significant effect of pellet availability (two-way ANOVA/Bonferroni multiple-comparison test; stim, p = 0.1094; F(1,4) = 4.211; pellet, **p = 0.016; F(1,4) = 58.44; interaction, p = 0.1146; F(1,4) = 4.045). C. Preference index calculation for females’ and males’ preference for the Pellet+Stim port over other rewarded ports for mice receiving MCH optogenetic stimulation in LH. No significant difference was observed (two-sample t-test, p=0.8313). D. Poke percentages on the final day of the StimON phase of the ORCA paradigm for females (top) and males (bottom) receiving MCH optogenetic stimulation in NAc. No significant differences between ports were observed. E. Cumulative poking behavior over the entire 12hr dark cycle on the final day of the StimON phase of the ORCA paradigm for females (top) and males (bottom) receiving MCH optogenetic stimulation in NAc. No significant differences between ports were observed. F. Preference index calculation for females’ and males’ preference for the Pellet+Stim port over other rewarded ports for mice receiving MCH optogenetic stimulation in NAc. No significant difference was observed (two- sample t-test, p=0.0824).

Females receiving optogenetic stimulation in NAc (Figure 7D, top) showed a significant effect of pellet availability (two-way ANOVA/Bonferroni multiple-comparison test; stim, p = 0.1856; F(1,10) = 2.021; pellet, ***p = 0.0001; F(1,10) = 35.00; interaction, p = 0.9888; F(1,10) = 0.0002062). Males receiving optogenetic stimulation in NAc (Figure 7D, bottom) showed a significant effect of pellet availability (two-way ANOVA/Bonferroni multiple-comparison test; stim, p = 0.1094; F(1,4) = 4.211; pellet, **p = 0.016; F(1,4) = 58.44; interaction, p = 0.1146; F(1,4) = 4.045). This preference emerged over the course of the 12 h of stimulation (Figure 7E). In our preference index measure, male mice receiving MCH optogenetic stimulation in NAc showed a higher, though not significant, preference index than their female counterparts (Figure 7F). Our data demonstrates that MCH has similar overall effects in males and females, but supports previous reports that the MCH system may more potently impact behavior in males.

## Discussion

The earliest investigations into the role of MCH demonstrated its ability to increase feeding. Knockout of the *Pmch* gene leads to reduced food consumption and an anorexia-like phenotype while overexpression of MCH results in increased feeding and body weight gain (Shimada et al., 1998; Ludwig et al., 2001).

Pharmacological evidence also points to a role for MCH peptide in food consumption; injection of MCH into the cerebral ventricles, and a number of other specific brain regions, results in increased food consumption (Rossi et al., 1997; Georgescu et al., 2005; Guesdon et al., 2009). However, more recent work using optogenetic and pharmacogenetic approaches do not have clear findings linking the MCH system to increased food consumption. Dilsiz et al. show that stimulation of MCH neurons with either pharmacogenetic or optogenetic approaches does not induce increased feeding; indeed, optogenetic activation could slightly but significantly reduce feeding (Dilsiz et al., 2019). We similarly do not observe increased consumption when optogenetically stimulating MCH neurons in the LH. While there has been interest in developing MCH receptor antagonists for treatment of obesity, it is worth noting that none have reached the clinic and they seem to have modest effect sizes at best (Potter and Burgess, 2022). A key difference between the majority of the early work in this area and the more recent application of specific circuit mapping tools is that recent experiments involve activation of the MCH ‘system’ (i.e., the neurons that express MCH) vs MCH peptide release or MCH receptor activation per se. Some functions of the MCH system are a result of glutamatergic, and potentially GABAergic, release from these neurons and it is unclear the extent to which MCH peptide itself is released in response to *in vivo* pharmacogenetic or optogenetic activation (Liu et al., 2022; Beekly et al., 2023). Furthermore, it is possible that glutamate, GABA, and/or other peptides released from MCH neurons may have differing or opposing effects to MCH peptide.

MCH peptide injections into the NAc increase food intake (Georgescu et al., 2005; Guesdon et al., 2009), while pharmacogenetic activation of the MCH neurons that project to the NAc can drive modest increases in feeding in male mice and no change in females (Terrill et al., 2020). Here, we do not see increased food intake with optogenetic stimulation of MCH terminals in the NAc, in neither a homecage environment nor in the optogenetics-reinforced consumption assay. There are several potential explanations for this apparent discrepancy. Previous work has been performed predominantly in rats, and there is evidence of differences in rat vs mouse MCH systems, at least at the anatomical level (Croizier et al., 2010).

Alternatively, optogenetic vs pharmacogenetic manipulation of MCH neurons may result in differential release and/or release patterns of glutamate vs MCH peptide, ultimately resulting in different postsynaptic effects. Those MCH neurons that project to the NAc likely also exhibit axon collaterals in several other brain regions given the widespread nature of MCH projections, whereas optogenetic stimulation of terminals is more likely to limit this activation to the NAc region (though there is evidence that in certain conditions optogenetic activation of terminals may result in antidromic stimulation of cell bodies (e.g., Jennings et al., 2013), and we cannot rule this out in the current study). It is also possible that our stimulation was not sufficiently strong to see the subtle feeding effects previously characterized. Our characterization of these mice showed clear light-evoked activation of MCH neurons in *ex vivo* slice experiments, but not all neurons were able to follow the 20 Hz light train, suggesting that transmitter release in the NAc may not have been maximal. We do demonstrate notable behavioral effects but it is possible that more sustained or higher frequency activation *in vivo* could have also driven a feeding effect. We refer to the effects of MCH on ‘non-homeostatic’ feeding here because we did not see changes in total food consumption with MCH stimulation (Figure 3B,C; Extended figure 4-1A) but did see reinforcing effects on consumption of food paired with MCH→NAc stimulation (Figure 4). It is indeed possible that the MCH system could drive both homeostatic and non-homeostatic consumption based on the prior literature and the caveats above.

A previous study demonstrated that MCH-KO mice do overeat in response to food restriction, a homeostatic motivator, but do not overeat in response to conditioned food cues (Sherwood et al., 2015). Our findings seem to complement this work, as we demonstrate an increase in preference for a specific food paired with MCH → NAc terminal stimulation, but not an increase in overall food consumption, suggesting that MCH terminal stimulation in the NAc may serve to reinforce consumption of food paired. The NAc has a well-established role in reward and reinforcement learning, particularly as it relates to mesolimbic dopaminergic inputs (Salgado and Kaplitt, 2015; Sackett et al., 2017; Castro and Bruchas, 2019). The MCH system has been implicated in regulation of dopamine activity, suggesting that this interaction may underlie reinforcing effects of MCH in the NAc (Sears et al., 2010; Domingos et al., 2013; Chee et al., 2019; Duncan Spencer et al., 2024). A recent study demonstrated that MCH in the ventral tegmental area modulates both midbrain dopaminergic and glutamatergic neurons (Duncan Spencer et al., 2024), but there is also evidence of more local interactions between MCH neurons and dopamine release in the NAc. MCH peptide has been shown to enhance the postsynaptic effects of the dopamine receptor agonist apomorphine in the NAc, and to increase NAc neuron firing when both D1R and D2R are activated (Chung et al., 2009; Hopf et al., 2013); though MCH peptide itself, as opposed to neural activation, likely decreases dopamine release in the NAc and the activity of NAc neurons (Sears et al., 2010; Chee et al., 2019). As noted above, MCH neurons release multiple neurotransmitters and peptides in addition to MCH (Beekly et al., 2023). Previous work suggests that MCH neurons mediate the post-ingestive value of sucrose consumption; this work also demonstrates that activation of MCH neurons can increase dopamine release in the NAc (Domingos et al., 2013). Follow up work shows that sucrose preference is mediated by glutamatergic release from MCH neurons, not the MCH peptide (Schneeberger et al., 2018).

These studies suggest that the MCH system may act to enhance dopaminergic signaling or postsynaptic neural activity in the NAc, facilitating the reinforcement behavior observed here. However, they also indicate that this effect might not be mediated by MCH peptide release.

We also demonstrate that, while stimulation of MCH terminals in the NAc is reinforcing, it does not increase food pellet consumption or increase REM sleep. This is despite MCH neuron activation in the LH increasing REM sleep as we show here and as others have previously demonstrated (Jego et al., 2013; Varin et al., 2018; Kroeger et al., 2019). Relatively few studies of the MCH system have investigated more than one MCH-related behavior, or the role of efferent projections, despite there being evidence of subpopulations of MCH neurons, both at the level of anatomy and gene expression (Cvetkovic et al., 2003, 2004; Beekly et al., 2023; Miller et al., 2024). At least two subtypes of MCH neurons have been characterized based predominantly on co-expression of MCH and Cocaine and amphetamine regulated transcript (CART), with differing projection and electrophysiological profiles (Cvetkovic et al., 2004; Fujita et al., 2021; Miller et al., 2024), but it is unclear how they differentially impact physiology and behavior. Our data also point to distinct subpopulations of MCH neurons based on anatomical projection profile. Our demonstration that MCH → NAc terminal stimulation does not result in the same REM sleep effect as LH MCH cell body stimulation is suggestive of at least two populations of MCH neurons, one of which drives REM sleep while the other enhances the reward value of consumption. While our results help to understand how the MCH system can regulate many disparate behaviors, including food consumption, more work is needed to fully elucidate the structure-function relationship of the MCH system.

## Conflict of Interest statement

The authors declare no competing financial interests.

## Acknowledgements

We thank Dr. Shelly Flagel, Dr. Liam Potter, Elizabeth Rose Burgess, and all members of the Burgess lab for helpful discussion. Special thanks to Lucia Kim for assisting with early versions of these experiments. This work was supported by 1F31DK135283-01(KLF), NIDA T32 DA7281(KLF), a Michigan Diabetes Research Center Pilot and Feasibility Award, a Whitehall Foundation new investigator grant, and 1R01DK129366-01 (CRB).

## Author Contributions

KLF and CRB contributed to project conceptualization. KLF contributed to data curation.

KLF, HCL, LB, and CRB contributed to formal data analysis. KLF and CRB contributed to funding acquisition.

KLF, HCL, LB, TC, JRE, J Manna, LZ, and J Mattis contributed to the research investigation process, i.e. performing experiments and collecting data.

KLF, LB, TC, LZ, J Mattis and CRB contributed to development and design of methodology. KLF, LB and J Manna and CRB were responsible for project administration and management.

KLF, JRE and CRB contributed to programming, software development, and implementation of both acquisition and analysis code.

TC, J Manna and CRB contributed to the provision of resources including study materials, reagents, materials, animals.

CRB was responsible for oversight and leadership supervision. KLF, LB, HCL and CRB contributed to data visualization.

KLF and CRB wrote the original draft of the manuscript.

All authors contributed to reviewing and editing the manuscript.

## Extended Figure Legends

**Extended Data Figure 3-1. Optogenetic stimulation of the MCH system did not change feeding behavior in control mice.**

A. Schematics showing fiber placements in the LH and NAc.

B. Control mice with fiber in LH show no significant change in pellet consumption (top) REM bout length (middle) or REM bout number (bottom) with light stimulation regardless of fiber location (paired t-test, p>0.05). Pink and blue dots denote female and male mice, respectively.

C. Control mice with fiber in NAc show no significant change in pellet consumption (top) REM bout length (middle) or REM bout number (bottom) with light stimulation regardless of fiber location (paired t-test, p>0.05). Pink and blue dots denote female and male mice, respectively.

D. Control mice with fiber locations combined show no significant change in pellet consumption (top) REM bout length (middle) or REM bout number (bottom) with light stimulation regardless of fiber location (paired t-test, p>0.05). Pink and blue dots denote female and male mice, respectively.

**Extended Data Figure 3-2. Feeding architecture was not altered as a result of MCH optogenetic stimulation.**

A. Feeding architecture measurements for mice receiving MCH optogenetic stimulation. “Feeding bouts” were calculated by specifying an interbout-interval of 2 minutes. Pellet consumption events that were more than 2 minutes apart were separated into separate “bouts” of feeding, while those closer than 2 mins together were considered part of the same “bout.” Number of feeding bouts did not significantly change with optogenetic stimulation in LH (paired t-test, p>0.05). Pink and blue dots denote female and male mice, respectively.

B. Number of pellets per feeding bout did not significantly change with optogenetic stimulation in LH (paired t-test, p>0.05). Pink and blue dots denote female and male mice, respectively.

C. Number of feeding bouts did not significantly change with optogenetic stimulation in NAc (paired t-test, p>0.05). Pink and blue dots denote female and male mice, respectively.

D. Number of pellets per feeding bout did not significantly change with optogenetic stimulation in NAc (paired t-test, p>0.05). Pink and blue dots denote female and male mice, respectively.

**Extended Data Figure 3-3. Optogenetic stimulation of MCH neurons did not produce sex-specific differences in REM sleep or pellet consumption.**

A. There were no sex-specific differences in REM bout length (left), number of REM bouts (middle) or pellets consumed (right) with MCH optogenetic stimulation in LH (paired t-test, p>0.05).

B. There were no sex-specific differences in REM bout length (left), number of REM bouts (middle) or pellets consumed (right) with MCH optogenetic stimulation in NAc (paired t-test, p>0.05).

**Extended Data Figure 4-1. Optogenetic stimulation did not significantly impact feeding behavior.**

A. Optogenetic stimulation of MCH neurons in the LH or MCH terminals in the NAc did not increase the number of pellets consumed during the optogenetics reinforced consumption assay. Pink and blue dots denote female and male mice, respectively.

B-C. There was no difference in feeding architecture measurements for mice receiving optogenetic stimulation. Pink and blue dots denote female and male mice, respectively.

**Extended Data Figure 4-2. Optogenetic stimulation did not cause a preference for stim-paired ports in control mice.**

A. Control mice with optic fibers implanted in the LH showed a non-significant preference for pellet-paired ports in the StimON phase. Pink and blue dots denote female and male mice, respectively.

B. Control mice with optic fibers implanted in the NAc showed a non-significant preference for pellet-paired ports in the StimON phase. Pink and blue dots denote female and male mice, respectively.

## Notes

### Competing Interest Statement

The authors have declared no competing interest.

