## Extended Data Figures for "Melanin concentrating hormone projections to the nucleus accumbens enhance the reward value of food consumption and do not induce feeding or REM sleep"

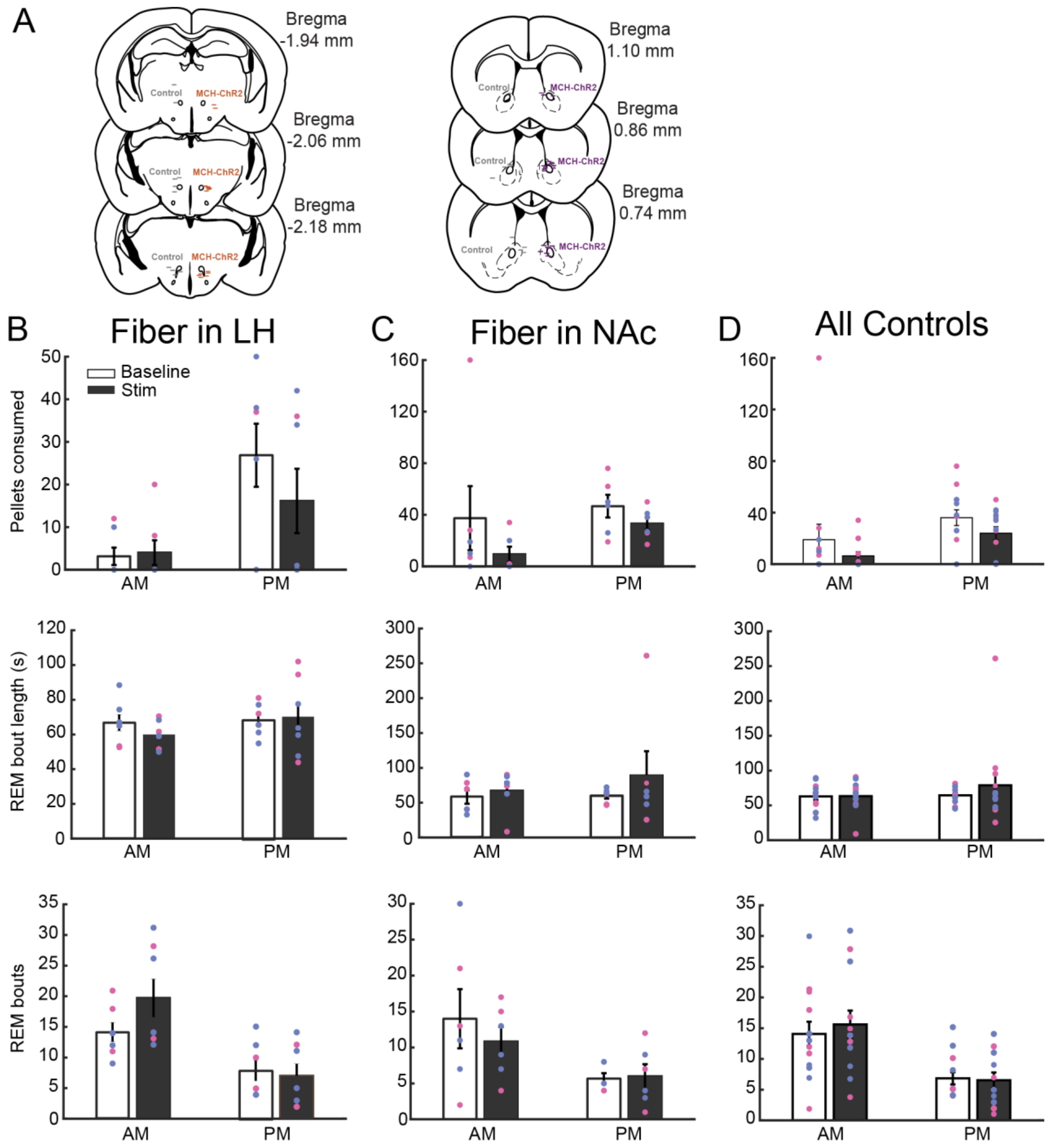


**Extended Data Figure 3-1. Optogenetic stimulation of the MCH system did not change feeding behavior in control mice.**

A. Schematics showing fiber placements in the LH and NAc.

B. Control mice with fiber in LH show no significant change in pellet consumption (top) REM bout length (middle) or REM bout number (bottom) with light stimulation regardless of fiber location (paired t-test, $p>0.05$). Pink and blue dots denote female and male mice, respectively.

C. Control mice with fiber in NAc show no significant change in pellet consumption (top) REM bout length (middle) or REM bout number (bottom) with light stimulation regardless of fiber location (paired t-test, $p>0.05$). Pink and blue dots denote female and male mice, respectively.

D. Control mice with fiber locations combined show no significant change in pellet consumption (top) REM bout length (middle) or REM bout number (bottom) with light stimulation regardless of fiber location (paired t-test, $p>0.05$). Pink and blue dots denote female and male mice, respectively.

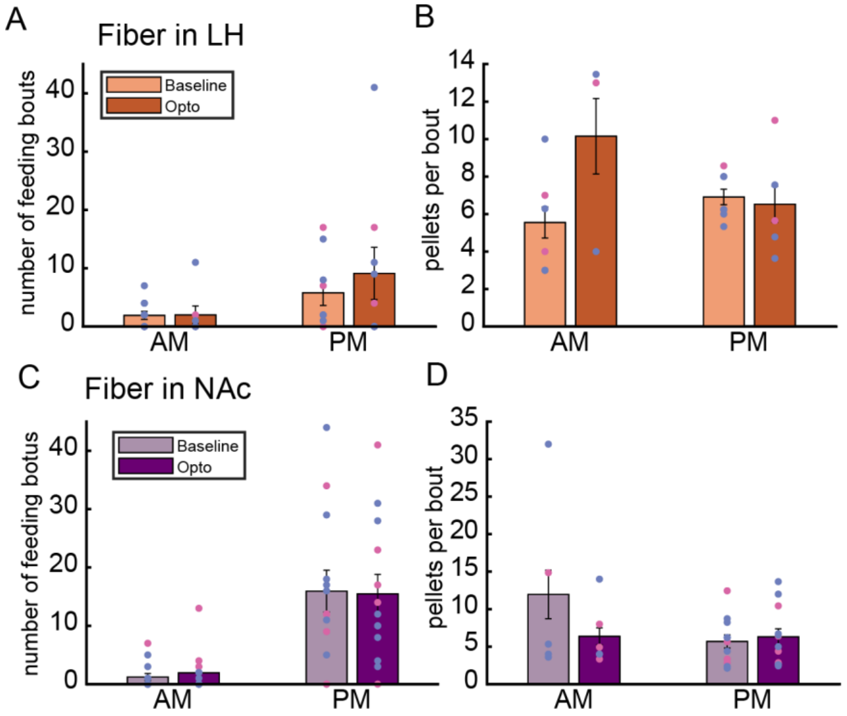


**Extended Data Figure 3-2. Feeding architecture was not altered as a result of MCH optogenetic stimulation.**

A. Feeding architecture measurements for mice receiving MCH optogenetic stimulation. “Feeding bouts” were calculated by specifying an interbout-interval of 2 minutes. Pellet consumption events that were more than 2 minutes apart were separated into separate “bouts” of feeding, while those closer than 2 mins together were considered part of the same “bout.” Number of feeding bouts did not significantly change with optogenetic stimulation in LH (paired t-test, $p>0.05$). Pink and blue dots denote female and male mice, respectively.

B. Number of pellets per feeding bout did not significantly change with optogenetic stimulation in LH (paired t-test, $p>0.05$). Pink and blue dots denote female and male mice, respectively.

C. Number of feeding bouts did not significantly change with optogenetic stimulation in NAc (paired t-test, $p>0.05$). Pink and blue dots denote female and male mice, respectively.

D. Number of pellets per feeding bout did not significantly change with optogenetic stimulation in NAc (paired t-test, $p>0.05$). Pink and blue dots denote female and male mice, respectively.

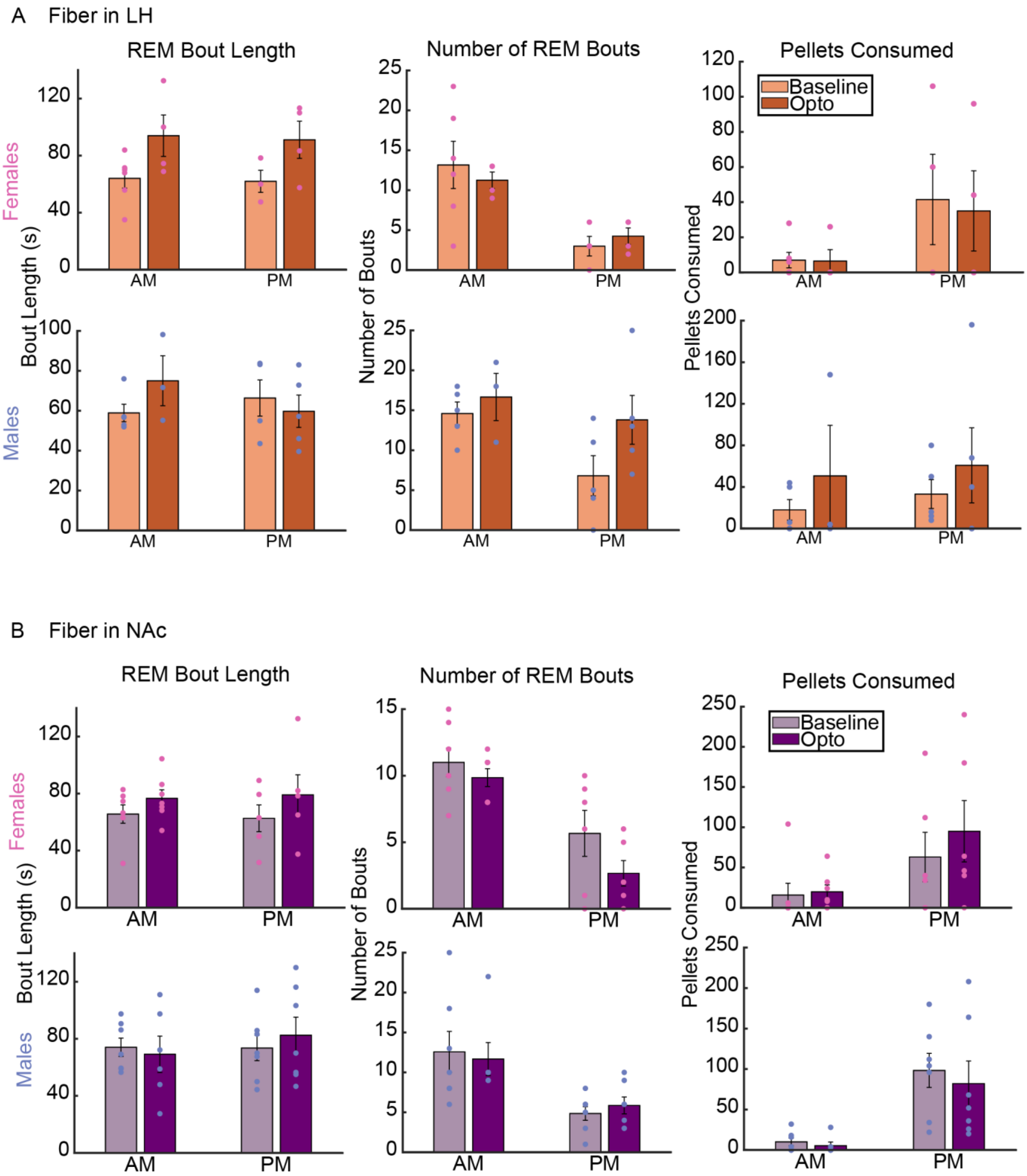


**Extended Data Figure 3-3. Optogenetic stimulation of MCH neurons did not produce sex-specific differences in REM sleep or pellet consumption.**

A. There were no sex-specific differences in REM bout length (left), number of REM bouts (middle) or pellets consumed (right) with MCH optogenetic stimulation in LH (paired t-test, $p>0.05$).

B. There were no sex-specific differences in REM bout length (left), number of REM bouts (middle) or pellets consumed (right) with MCH optogenetic stimulation in NAc (paired t-test, $p>0.05$).

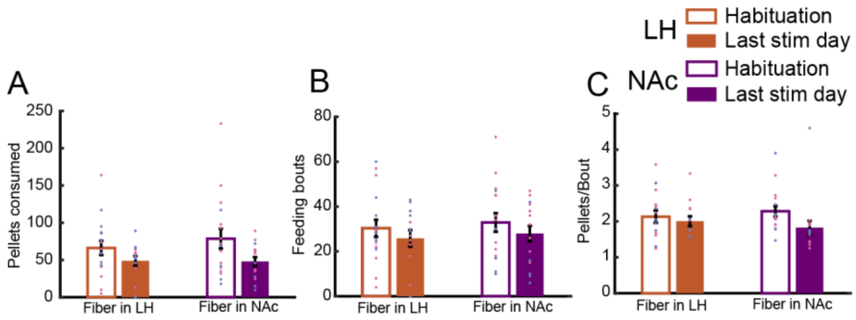


**Extended Data Figure 4-1. Optogenetic stimulation did not significantly impact feeding behavior.**

A. Optogenetic stimulation of MCH neurons in the LH or MCH terminals in the NAc did not increase the number of pellets consumed during the optogenetics reinforced consumption assay. Pink and blue dots denote female and male mice, respectively.

B-C. There was no difference in feeding architecture measurements for mice receiving optogenetic stimulation. Pink and blue dots denote female and male mice, respectively.

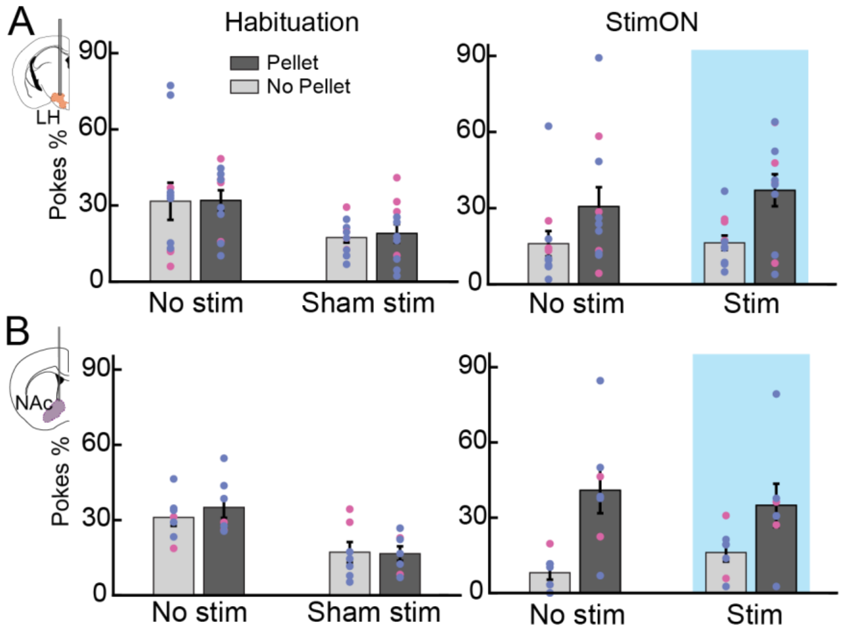


**Extended Data Figure 4-2. Optogenetic stimulation did not cause a preference for stim-paired ports in control mice.**

A. Control mice with optic fibers implanted in the LH showed a non-significant preference for pellet-paired ports in the StimON phase. Pink and blue dots denote female and male mice, respectively.

B. Control mice with optic fibers implanted in the NAc showed a non-significant preference for pellet-paired ports in the StimON phase. Pink and blue dots denote female and male mice, respectively.
